# Transgenerational exposure to deoxygenation and warming disrupts mate-detection in *Gammarus locusta*

**DOI:** 10.1101/2023.02.21.529266

**Authors:** Beatriz P. Pereira, Simon Neff, Francisco O. Borges, Eve Otjacques, Guilherme Barreto, Maddalena Ranucci, Mélanie Court, Rui Rosa, Tiago Repolho, José Ricardo Paula

**Affiliations:** MARE - Marine and Environmental Sciences Centre & ARNET - Aquatic Research Network, Laboratório Marítimo da Guia, Faculdade de Ciências da Universidade de Lisboa, Av. Nossa Senhora do Cabo, 939, 2750-374 Cascais, Portugal; Department of Biology, Faculty of Mathematics and Natural Sciences, University of Cologne, Zülpicher Straße 47b, 50674 Cologne, Germany; Carnegie Institution for Science, Division of Biosphere Sciences and Engineering, Church Laboratory, California Institute of Technology, 1200 E. California Blvd, Pasadena, CA 91125, USA; Departamento de Biologia Animal, Faculdade de Ciências, Universidade de Lisboa, Campo Grande, 1749-016 Lisboa, Portugal; Hawai‘i Institute of Marine Biology, School of Ocean and Earth Science and Technology, University of Hawai‘i, 46-007 Lilipuna Road, Kaneohe, HI 96744, USA

**Keywords:** ecology, climate change, reproductive behaviour, amphipod, gammarid, carry-over effects

## Abstract

Ocean deoxygenation and warming have been shown to pose a growing threat to the health of marine organisms and ecosystems. Yet, the potential for acclimation and adaptation to these threats remains poorly understood. The aim of this study was to evaluate the effects of transgenerational exposure to reduced oxygen availability and elevated seawater temperature on the chemosensory-dependent mating mechanisms of male amphipods *Gammarus locusta*. After exposure, the number of individuals that reached adulthood (in F0 and F2) was gauged, and adult males from F0 and F1 were subjected to behavioral trials to assess their capacity of long-distance female cue detection through quantification of (i) response time; (ii) first direction of movement; (iii) activity rate and (iv) proportion of time spent in female scent cues. Ocean warming induced mortality (especially in F2), and reduced oxygen availability had adverse effects on each of the investigated behavioral traits, which were amplified when combined with elevated temperature. Still, when compared to F0, the F1 generation demonstrated more adaptability (i.e., higher activity rate and preference for female odors) to the combination of the two stressors, suggesting positive carry-over effects. Nevertheless, full recovery to control levels was not observed. Altogether, this study indicates that future scenarios of ocean deoxygenation and warming have the potential to disrupt chemosensory-dependent mate-detection in amphipods, but also suggests possible behavioral adaptations. We call for greater research efforts on long-term impacts of ocean change on the behavioral and physiological processes of benthic coastal communities.

## 1. Introduction

The oceans play a vital role in maintaining the Earth’s climate by absorbing approximately 90% of the excess energy from greenhouse gases and up to 30% of the CO_2_ released into the atmosphere (Bindoff et al., 2019; Collins et al., 2019; Oschlies et al. 2018). The increased transfer of atmospheric gases, energy, and heat have led to changes in oceanic physicochemical processes (Bindoff et al., 2019). Long-term monitoring studies have found a significant rise in sea surface temperatures (SST), especially in coastal areas, where the temperature increase has been reported as +0.17°C per decade (Liao et al., 2015). Moreover, ocean warming (OW) together with increased nutrient discharge are key contributors to another major climate change-related phenomenon: ocean deoxygenation (OD). Rising temperatures promote oxygen loss by decreasing oxygen solubility and promoting stratification. Decreased oxygen mixing, increased oxygen outgassing (projected to increase from 1.6 to 4.3 gigatons. year^−1^ over the 21^st^ century), and decreased nutrient input, limiting photosynthesis and oxygen generation (Oschlies et al., 2018; Keeling et al., 2010; Laffoley and Baxter, 2019; Li et al., 2020). Compared to 1990s reference values, oxygen concentration in the ocean is predicted to decrease between 8 and 40 mmol.m^−3^ (Bindoff et al., 2019; Collins et al., 2019; Li et al., 2020).

OW and OD can affect marine organisms’ growth and reproduction, physiological stress, habitat compression, abundance, distribution, life cycles, and ecological interactions (Vaquer-Sunyer and Duarte, 2008). Although stressful conditions can inflict negative impacts on several species, they can also provide opportunities for others to adapt and even thrive under these changes (Munday et al., 2013; Reusch, 2014; Gallo and Levin, 2016). Exposure of a parental generation to stress drivers can lead to transgenerational acclimation, that is, the transmission of nongenetic traits to offspring (i.e., carry-over effects), which can positively influence the performance of subsequent generations under the same conditions (Sunday et al., 2014; Griffith and Gobler, 2017). However, this is not always the case, with some studies revealing negative carry-over effects which render the offspring more sensitive to stressors (Byrne, 2011; Griffith and Goble, 2017; Schade et al., 2014).

Many crustaceans dwell in coastal waters, frequently experiencing environmental fluctuations, with seawater physiochemical conditions varying on a seasonal, monthly, daily, or even hourly basis (Hofmann et al., 2011, Duarte et al., 2013). One would think that species that are already to substantial environmental fluctuations are likely to be relatively unaffected by future anthropogenic changes. Alternatively, these species may be living at or near their physiological limitations and thus have limited ability to cope with additional environmental changes (Hofmann et al., 2011, Baumann et al., 2015). This is especially true when multiple environmental drivers cause change at the same time, resulting in additive or synergistic impacts (Boyd et al., 2018). Overall, most studies focused on researching the effects of ocean warming and acidification, while deoxygenation and its interaction with other stress factors has been neglected (Borges et al., 2022).

Amphipods comprise an order of Crustacea and are particularly popular in ecological studies due to their ecological relevance, responsiveness to environmental stressors, and good suitability for breeding and experimentation in captivity (Reish 1993; Neuparth et al., 2002; van Riel et al., 2007; Lavaniegos and Ohman, 2021). Like other crustaceans, the reproductive success of these animals is influenced by their ability to detect chemical and olfactory stimuli during the search for suitable mates (Breithaupt and Thiel, 2010). The marine euryhaline species, *G. locusta* is widely distributed along the Atlantic coast of Europe, playing a key ecological role in coastal food webs (Costa and Costa, 2000). Mate-attraction in *G.locusta* is mediated by long-distance female pheromones, which trigger active searching by males, and by body-contact pheromones in the female exoskeleton, which causes males to engage in the so-called pre-copulative mate-guarding (Borowsky 1985; Borowsky and Borowsky 1987; Sutcliffe 1992). The potential disruption of its sexual behaviour and subsequent population decline can have far-reaching consequences for other trophic levels and, as a result, the health of entire coastal communities.

The present study aims, for the first time, to elucidate the potential transgenerational effects of OD and OW on the olfactory mate-detection and attraction of the amphipod *Gammarus locusta*, over two generations (F0 and F1). Understanding how reproductive traits may be affected by environmental change and whether these effects persist across generations may allow inferring possible changes in mating behaviour through physiological responses and thus the natural adaptability of *G. locusta* in warmer and less oxygenated waters.

## 2. Material and Methods

### 2.1. Collection and stock culture of amphipods

Wild *Gammarus locusta* were collected on July 2021 at the southern edge of the Sado Estuary, Portugal (38° 27′ N, 8° 43′ W) and transported to the aquatic facilities of Laboratório Marítimo da Guia (Cascais, Portugal). Organisms were laboratory cultured for two months within a semiclosed aquatic life support system (LSS) supplied with natural seawater. Culture conditions were as follows: seawater temperature 18.8°C ± 0.5 pH 7.97 ± 0.09; salinity 34.5 ± 0.6; dissolved oxygen 110.2% ± 2.5 (air saturation). Once a week, *G. locusta* individuals were sieved and separated according to size/age (i.e., newborns, juveniles, sub-adults, and adults). The gammarids were fed ad libitum with the green algae *Ulva sp*.

### 2.2. Transgenerational stress exposure

The experimental setup followed a full factorial design manipulating seawater temperature and oxygen levels. Two generations of *G. locusta* were exposed to a total of four treatments: i) present-day scenario (control; 18.1 ± 0.13°C, 101.6 ± 0,95% air saturation, 7.78± 0.16 mg O_2_ L^−1^); ii) warming scenario (W; 21 ± 0.22°C, 101.5 ± 0.95% air saturation, 7.37±0,08 mg O_2_ L^−1^); iii); deoxygenation (D; 18 ± 0.19°C, 90.7±2.5% air saturation, 7.05±0.47 mg O_2_ L^−1^); iv) warming + deoxygenation (WD; 21 ± 0.41°C, 90,6 ± 2,62 % air saturation, 6.58±0.13 mg O_2_ L^−1^). pH and salinity values were kept at 8.00 ± 0.04 and 35 ± 0.4, respectively. Dissolved oxygen (DO) concentration was adjusted automatically using an Arduino controlling system (Mucha, 2020; hysteresis set at 0.2 mg O_2_ L^−1^) via solenoid valves connected to optical oxygen sensors (PyroScience FireStingO_2_, accuracy ± 0.1 mg O_2_ L^−1^). Solenoid valves injected certified N2 gas (Air Liquide, Portugal) into cylindrical mixing tanks, which continuously directed oxygen-limited seawater to the water baths via a 35-W water pump (TMC, V2 Power Pump, 2150 L h^−1^). The seawater temperature was controlled using seawater chillers (Frimar, Fernando Ribeiro Lda., Portugal) and submerged heaters (300W, TMC-Iberia, Portugal).

All treatment groups were supported by a flow-through LSS system that ensured a steady renewal of seawater. For each treatment, a glass tank served as a seawater bath for four 4-L replicate tanks (4 replicates per treatment, 16 in total). To simulate the natural environment, a thin layer of sand (~ 1 cm), rocks to provide shelter, and *Ulva* spp. (unrestrictedly available to avoid food privation and cannibalism) were added to each replicate. To complement the automatic systems, manual measurements of seawater parameters were conducted daily on each replicate tank: pH (pH meter VWR, pHenomenal, accuracy ± 0.005), temperature, and dissolved oxygen (oximeter VWR DO220, accuracy ± 0.3°C, ± 0.1 mg O_2_ L^−1^), and salinity (Hanna refractometer, accuracy ± 0.1 ppt). Additionally, total ammonia and nitrite levels were checked every three days using colorimetric tests (Tropic Marin, Germany). The mean values of physiochemical parameters for each treatment group are presented in Table S1 (Supplementary Material).

A total of 48 *G. locusta* mating couples, corresponding to the parental generation (F0), were used. These couples were randomly distributed across different treatment groups and replicates (24 couples per treatment and 6 couples per replicate). The organisms were acclimated for 7 days to the tanks before gradually exposed to changes in temperature (+1°C.day^−1^) and DO levels (−3%.day^−^ 1) until the set experimental values were reached.

During the exposure period, F0 gammarids were observed daily. Note that *G. locusta*’s reproductive behaviour includes a precopulatory guarding phase, during which the male holds and carries the female. Insemination occurs when the female molts and is ready to release eggs into the marsupium (Maranhao and Marques, 2003). As such, when couples separated and females were observed to be ovigerous, F0 males were allocated to small containers within their respective replicate tank and fed with *Ulva* spp. Similarly, ovigerous females were observed daily until the marsupium was found to be completely empty, at which point they were also separated in a second container. The same procedure was conducted for the F1 generation, approximately 30 days after birth. This was not carried out in the F2 generation, due to the high mortality observed in one of the treatments (WD). In this generation, survival to adulthood (30 days) was assessed.

Behavioral trials were carried out with adult male gammarids, which were randomly selected from small containers and starved for 24 hours prior to testing.

### 2.3. Binary-choice experiment

A total of 30 males from the F0 generation (nC= 10; nW = 7; nD = 6; nWD = 7) and 26 males from F1 generation (nC = 5; nW = 8; nD = 5; nWD = 8) were tested. A binary choice experiment, based on Borges et al. (2018b), was devised to investigate the transgenerational effects of increased temperature and/or lower oxygen levels on the male response to female scent signals (Figure S1, Supplementary Material). Two rectangular arenas (60 cm × 20 cm × 3 cm) were placed over two identical boxes on an evenly levelled surface (1&2, Figure S1a). Each of the arenas was connected to two 7-L cylindric header tanks (A-D, Figure S1a), placed on a high platform, which provided two opposing gravitational water masses, with distinct olfactory cues, to the arena via water inlets (A, Figure S1b) on each side: i) one containing female scent cues and ii) one containing clean filtered seawater (flow rate: 0.2 L m^−1^). These water masses were then drained at the center of the arena (B, Figure S1b), through a cylindric outlet (2 cm height) covered with a filter mesh (500 m), thus creating two immiscible water currents. Prior to the behavioral trials, preliminary tests were conducted to ensure optimal flow rates and avoid mixing of the two opposing flows, using blue food coloring and visual analysis as in Borges *et al*. (2018b).

For the preparation of the olfactory cues, two 200-L tanks were filled with seawater at two treatment temperatures (18°C & 21°C). Each tank served as a water bath to stabilize the temperature of six partially immersed 16-L containers with filtered and aerated seawater. To produce the test cues (i.e., female scent), female gammarids (n = 28) were taken at random from the stock culture and added to three containers at each temperature tank, where they remained for 48 h prior to behavioral trials to ensure pheromone accumulation. The other three containers in each temperature tank, with only filtered seawater, served as control cues. Gammarids were moved from the containers to the stock culture, and the resulting volumes were transferred to the header tanks, allowing for a total of 12 runs per testing day, after which a 48-hour interval was required to produce new test cues. Prior to each test run, the water parameters (i.e., temperature, DO, pH, and salinity) of the header tanks were measured to ensure that the different water masses shared the same conditions.

Each choice arena was filled with seawater at the appropriate temperature (i.e., respective to the male’s treatment) until it reached the center drain and a 2 cm water height. The behavioral trial started by placing a single male in a designated starting zone (in the center of the arena; C, Figure S1B), where it was held in a perforated container for 10 minutes of acclimation. The provided acclimation time was to allow the male to settle in the starting zone and ensure that both water masses reached the center of the arena, allowing the gammarid to sample both currents (Borges et al., 2018b). Following acclimation, the male was released from the container and allowed to swim freely throughout the arena (i.e., between association zones; D&E, Figure S1b) for another 10 min.

Two test runs were conducted simultaneously (two individuals, two different arenas; Figure S1a) in an isolated room and recorded from above with a video camera. Males from each treatment were chosen at random and tested twice (with cue input-side reversal). The connections between arenas and tanks were randomized and reassigned after each run, as was the filling of the tanks with either test or control cues. After each test run, the test subjects were carefully removed and the arenas (1&2; Figure S1) and the header tanks (A-D; Figure S1) were thoroughly rinsed, first with fresh water and then with salt water to remove any traces of test-cues and conspecific odors.

Video analysis was performed by a blind observer using BORIS 7.12.2 (Friard and Gamba, 2016). Four distinct traits of the male individuals were quantified: i) response time (i.e., the time required for a male gammarid to make its first initial movement toward one of the two association zones, after being released from the acclimation container); ii) the proportion of first choice (i.e., first movement directed towards female-scented association zone); iii) cumulative time (i.e., the proportion of time spent in each association zone); iv) activity rate (i.e., the proportion of time actively moving).

### 2.4. Statistical analyses

Data analysis of behavioral outputs was performed through generalized linear mixed models (GLMM) using the negative binomial (number of individuals, response time), binomial (first choice), and Gaussian (cumulative time & activity rate) residual distributions. All models used temperature, oxygen, and generation as categorical fixed factors, and replicate ID as random factors. Full models, with all possible interactions, were tested using the function ‘glmmTMB’ from the package ‘glmmTMB’ (Brooks et al., 2017) and the function ‘Anova’ from the package ‘car’ (Fox and Weisberg, 2011) in R, version 3.4.3 (RStudio Team, 2022). Post-hoc multiple comparisons were performed using the package ‘emmeans’ and applying Tukey corrections (Lenth, 2020). Model assumptions and performance were validated using the package “performance” (Lüdecke et al., 2021). Data exploration was conducted using the HighstatLibV10 R library from Highland Statistics (Zuur et al., 2009; - 2010).

## 3. Results

The number of gammarids that reached adulthood declined in the W treatment (Figure 1: χ^2^ = 6.967, d.f. = 1, p = 0.008) relative to the control between F1 and F2 (χ^2^ = 6.352, d.f. = 1, p = 0.012). Furthermore, a significant interacting effect was found between generation and temperature was found (χ^2^ = 9.184, d.f. = 1, p = 0.002).

**Figure 1:**
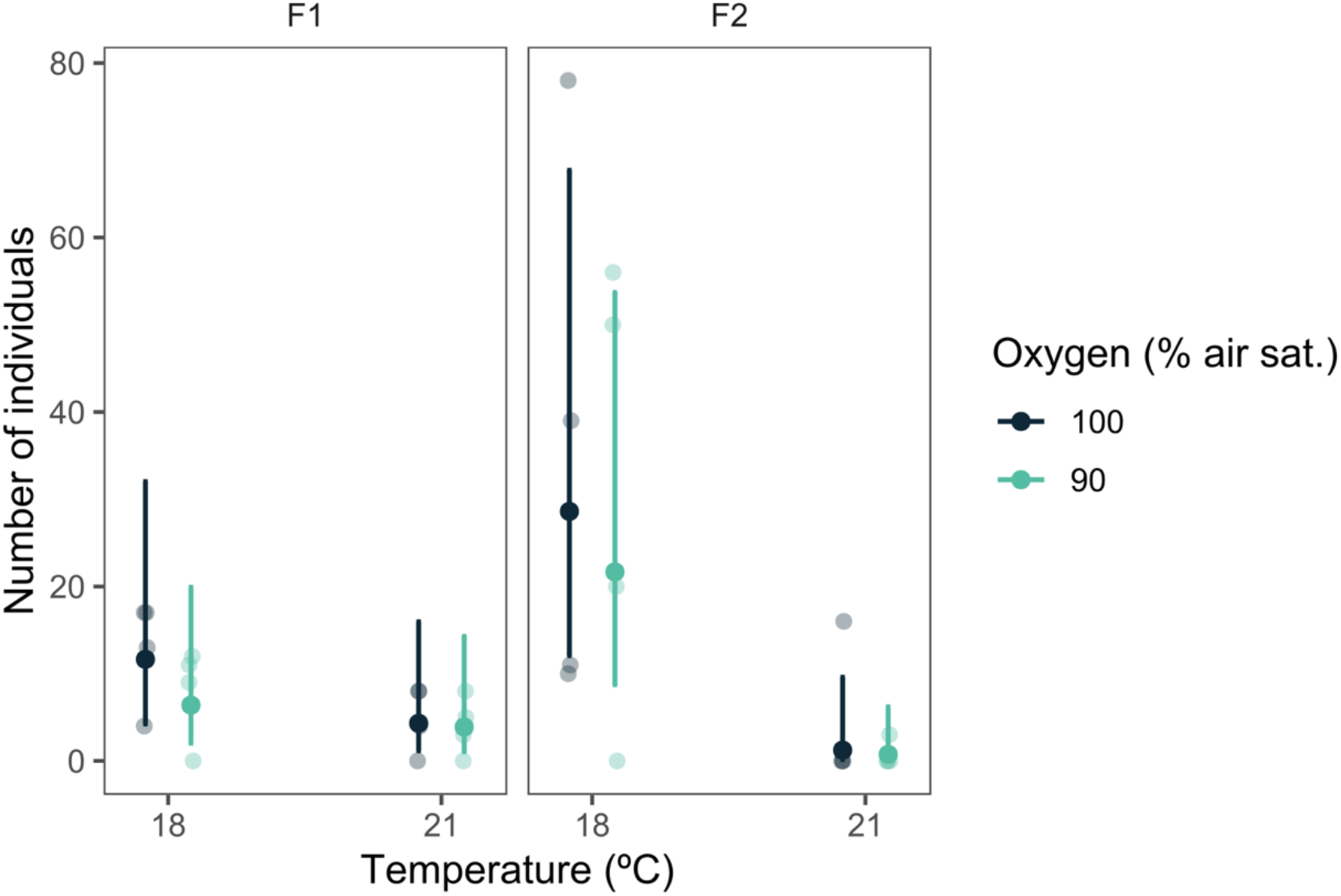
Number of individuals of *G. locusta* that reached adulthood in two successive generations (F1 & F2) exposed to four treatments with distinct temperatures (18 °C & 21 °C) and oxygen levels (90 % & 100 % air saturation). Back-transformed predicted means ± 95 % CI from the model and raw data values are presented.

There were no significant differences in the mean response time across generations or treatment groups or the interaction between generation and treatments (Figure 2; χ^2^ = 1.170, d.f. = 1, p = 0.279).

**Figure 2:**
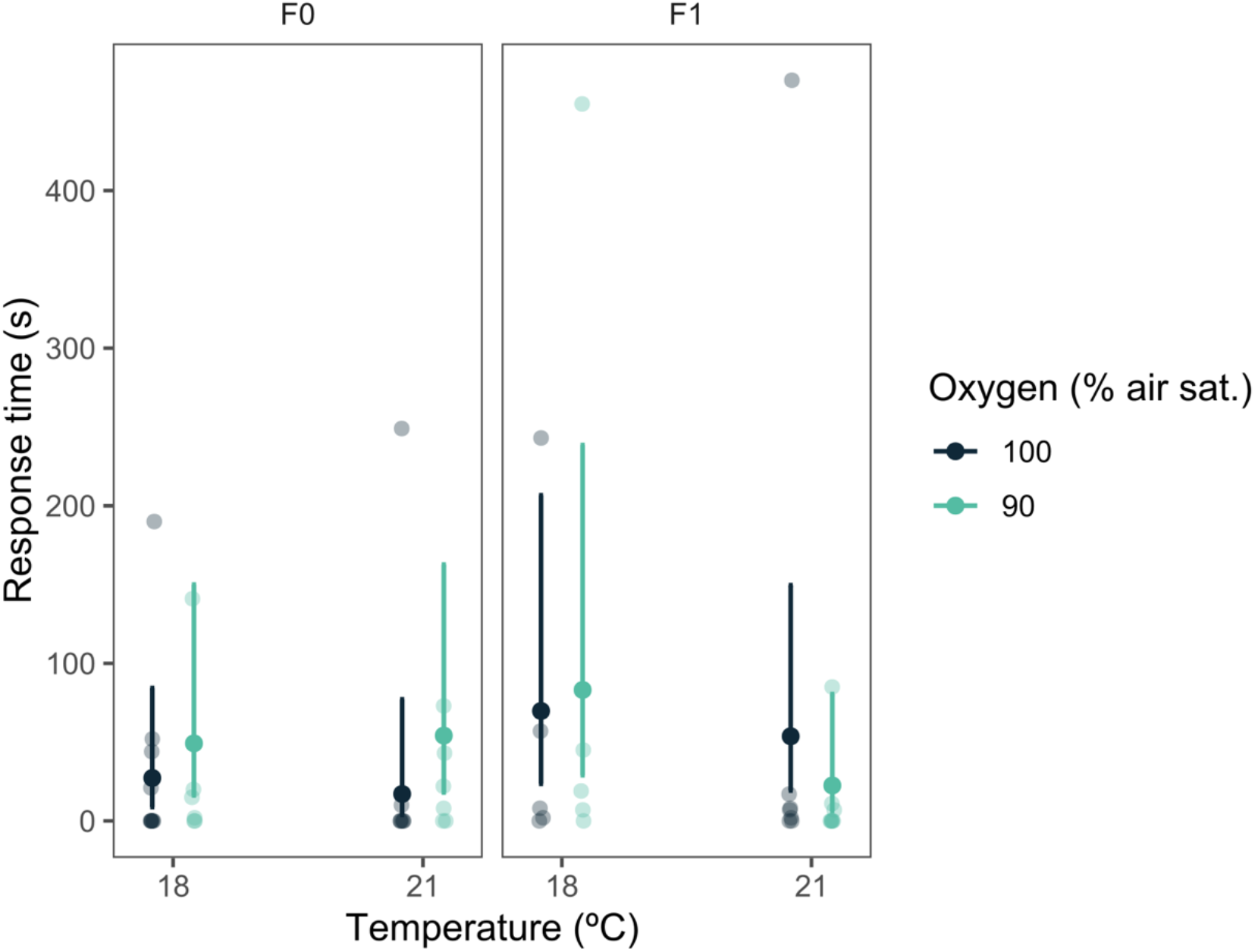
Response time (s) of male gammarids from two successive generations (F0 & F1) and four treatment groups varying in temperature (18 °C & 21 °C) and oxygen levels (90 % & 100 % air saturation). Back-transformed predicted means ± 95 % CI from the model and raw data values are presented.

Regarding the proportion of first choice for the female-scented association zone, no significant differences were detected across generations or treatment groups (Figure 3; χ^2^ = 0.748, d.f. = 1, p = 0.387).

**Figure 3:**
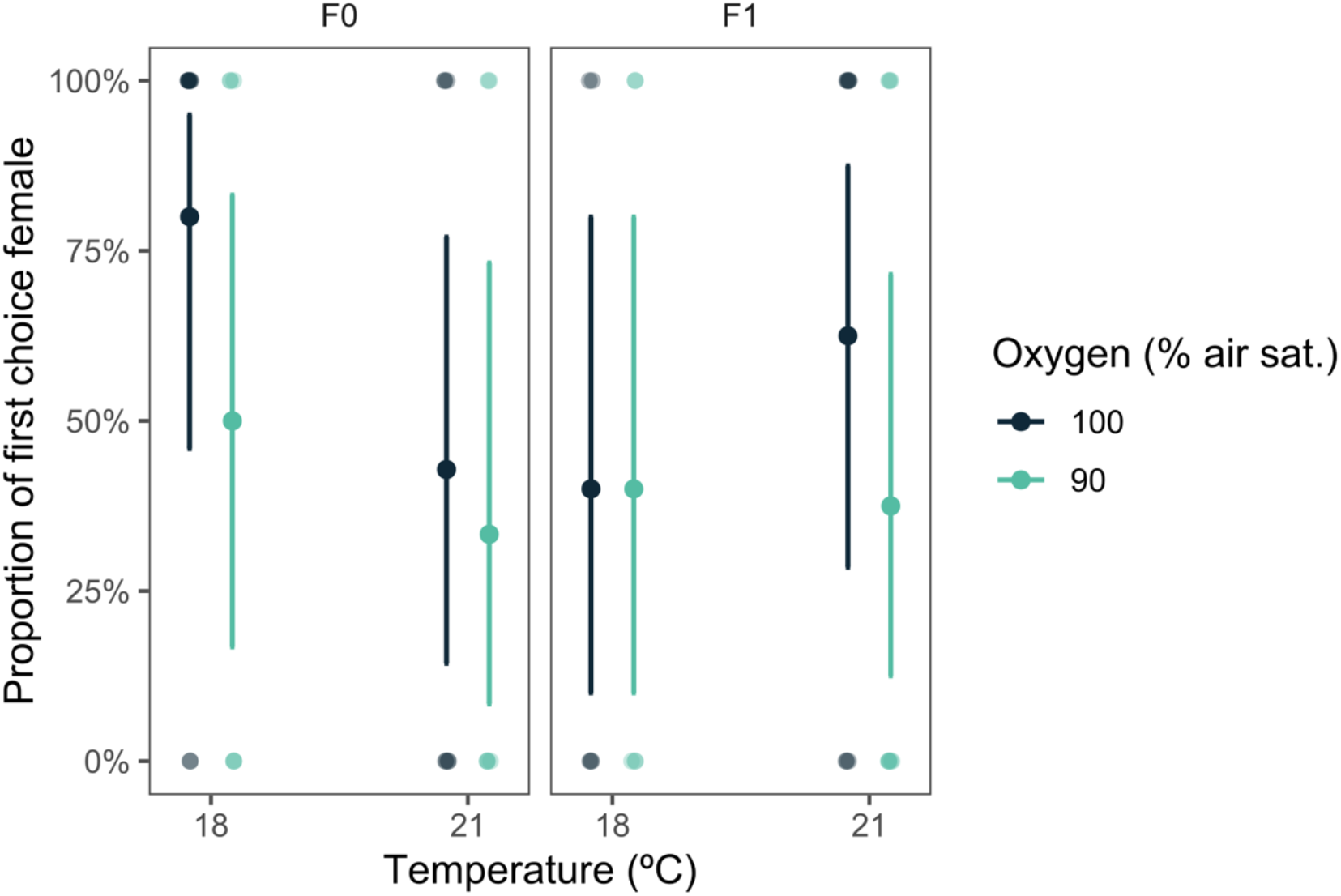
Proportion of first choices (%) of male gammarid from two successive generations (F0 & F1) and four treatment groups with different temperature (18 °C & 21 °C) and oxygen levels (90 % & 100 % air saturation). Back-transformed predicted means ± 95 % CI from the model and raw data values are presented.

However, when looking at the proportion of time spent by F0 gammarids in the association zone with the female cue, a significant reduction was observed in the WD treatment. On the other hand, the preference of F1 males for female scent returned to baseline levels in gammarids born in WD (Figure 4; χ^2^ = 7.705, d.f. = 1, p = 0.006).

**Figure 4:**
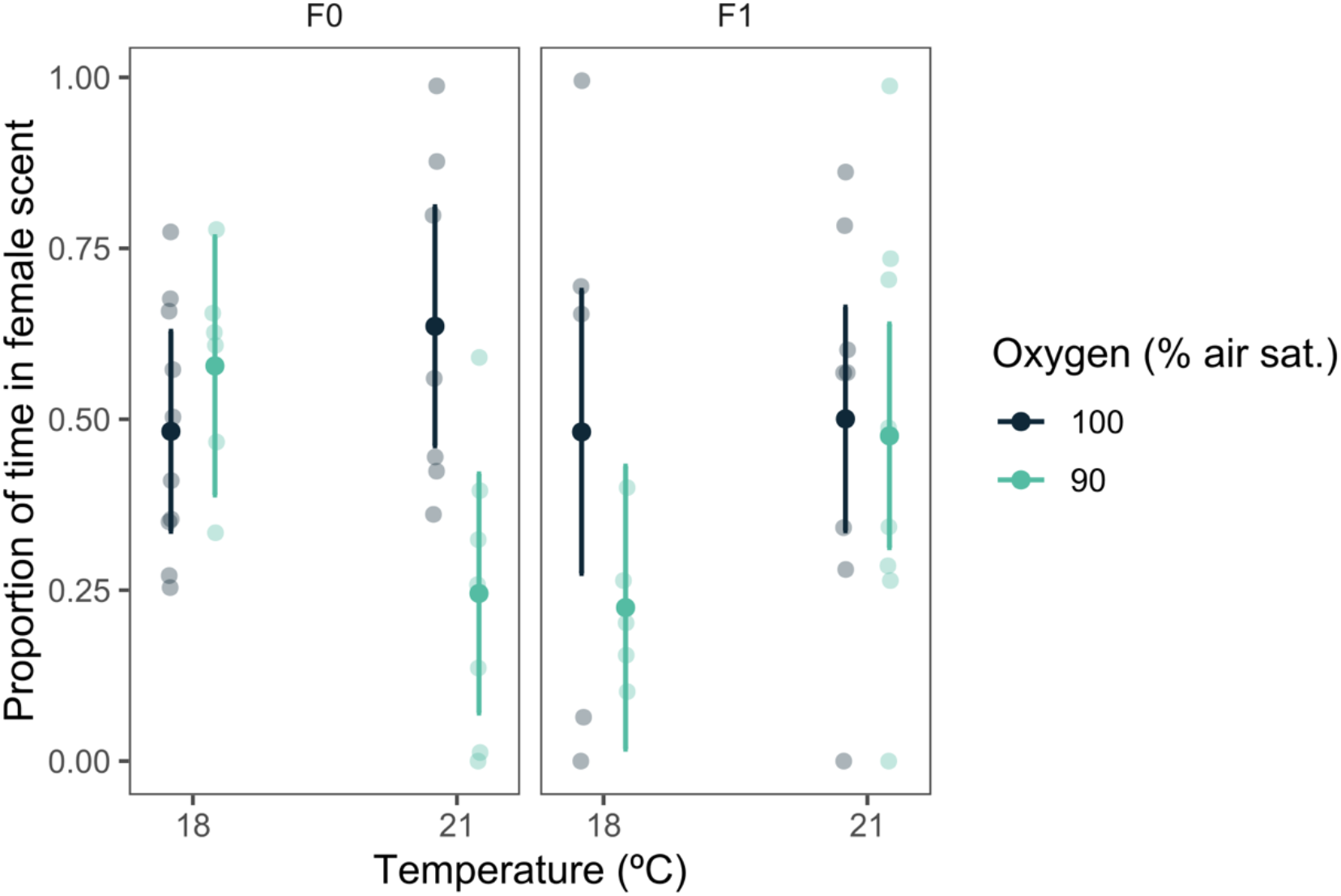
Proportion of time spent in the female-scented association zone by male gammarid from two successive generations (F0 & F1) and four treatment groups with different temperature (18 °C & 21 °C) and oxygen levels (90 % & 100 % air saturation). Back-transformed predicted means ± 95 % CI from the model and raw data values are presented.

Finally, looking at the activity rate of gammarids, the two treatment groups of F0 generation exposed to reduced DO levels were less active than the oxygen-saturated ones. However, in the subsequent generation F1, the opposite was observed, with males born in the D and DW treatments showing higher activity rates (Figure 5; χ^2^ = 4.571, d.f. = 1, p = 0.033) than those in C and W with 100% air saturation. The activity rates of the fully oxygenated treatments were not only surpassed by the deoxygenated groups in the F1 generation but also decreased compared to the F0 generation, especially in the C treatment group.

**Figure 5:**
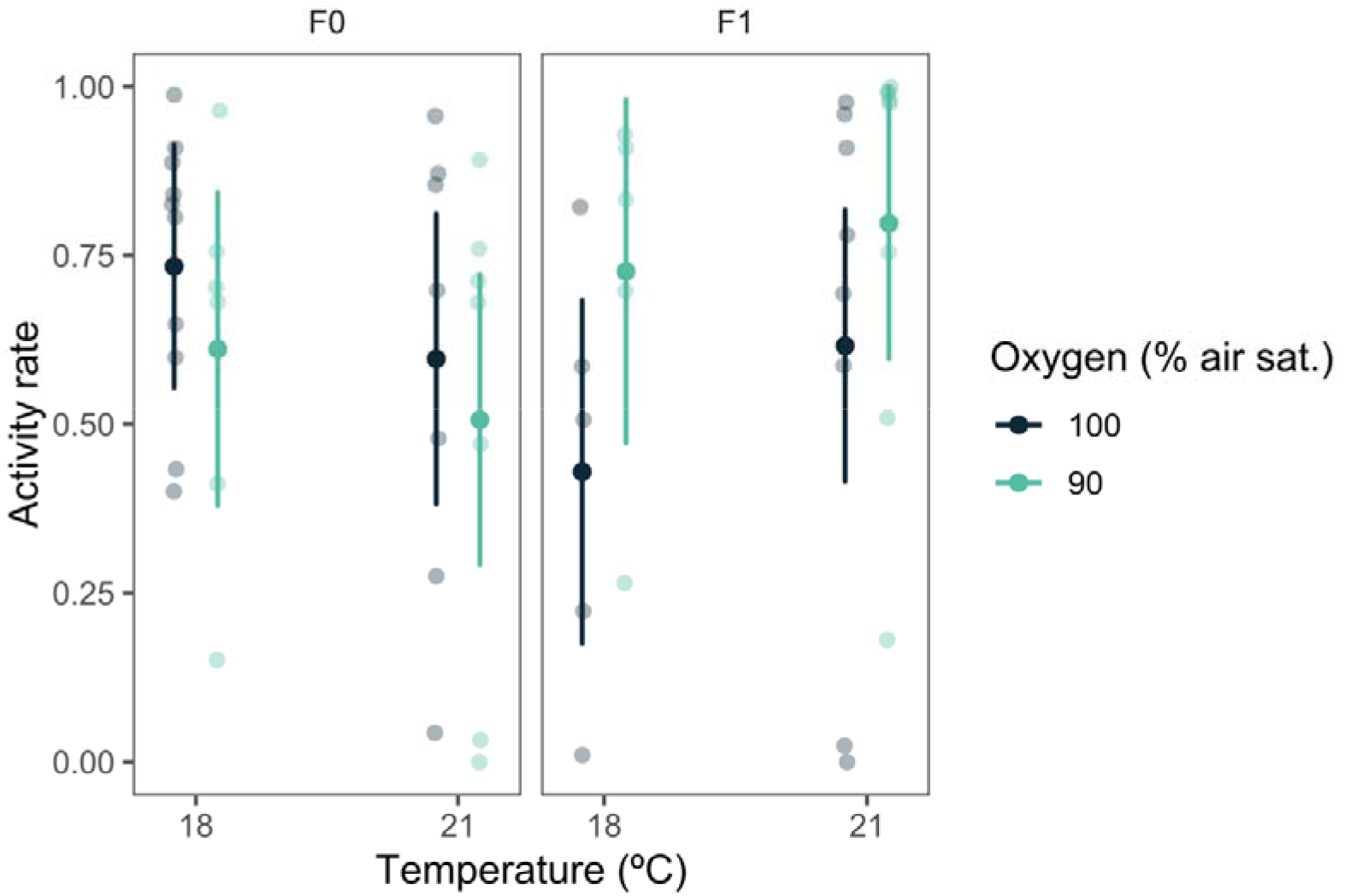
Activity rate of male gammarids from two successive generations (F0 & F1) exposed to four treatments with distinct temperatures (18 °C & 21 °C) and oxygen levels (90 % & 100 % air saturation) Back-transformed predicted means ± 95 % CI from the model and raw data values are presented.

## 4. Discussion

The results of this study indicate that transgenerational exposure of *Gammarus locusta* to ocean warming prevents the population growth expected in favorable conditions (laboratory rearing), as reflected in the growth observed in control conditions across generations (F0 - F2). This suggests that warming reduces fertility or results in direct mortality. The impacts of warming on survival have been previously documented, with mortality reaching 80% after only 21 days of exposure to + 4°C (Cardoso et al., 2018). However, mortality seems to escalate over generations, which is illustrated by the interaction between temperature and generation. This could indicate that ocean warming affects population growth, but only if the second filial generation is considered, denoting an inability of this species to adapt transgenerationally to increasing temperatures.

In contrast, both reduced oxygen availability and elevated water temperature can inpair the chemosensory-dependent mating mechanisms of male amphipods. Specifically, we observed that F0 male gammarids from WD treatment exhibited a significantly lower attraction for female pheromones compared to the other treatment groups (i.e., the proportion of time spent in the cue zone, Figure 4). However, following transgenerational acclimation, F1 males raised under WD conditions recovered their pheromone attraction to control levels, indicating that they were able to adapt to the adverse environmental conditions on a transgenerational level (i.e., positive carryover effects). Notably, although there was no difference in association zone preference between the single stressor groups (C, W, and D) for F0 generation males, F1 males exposed to low oxygen content showed a reduced proportion of the total trial time spent in the association area near the female cue. This indicates that chemosensory and olfactory perception in these animals was only affected by transgenerational oxygen deprivation. Since males exposed to the combined stressors (i.e., WD) were able to recover within one generation, this suggests that the synergistic effect of both stressors enhanced the protective carryover effects between generations. However, this behavioral adaptation was insufficient to mitigate the adverse effects on *G. locusta* physiology, as seen by the collapse of the F2 population. Thus, further research including physiological endpoints, such as monoamines or oxidative/nitrogen stress biomarkers, would be useful to uncover the physiological mechanisms responsible for this decline.

Furthermore, exposure to deteriorating conditions associated with reduced oxygen concentrations significantly affected activity rates. While the two deoxygenated groups (D and WD) in the parental generation (F0) had a lower activity rate than the other two treatment groups, this pattern was reversed in the subsequent generation (F1), with the two deoxygenated groups displaying noticeably higher activity rates than those with 100% air saturation (Figure 5). It is possible that this increased activity rate of F1 males correlates with the considerably decreased response time observed in WD treatment (Figure 2). Increased activity rates could be a sign of increased motivation to find a mate, which in turn could lead to a decreased response time as the males are more actively searching for potential partners. However, it is important to note that the correlation between the increased activity rate and the decreased response time is not necessarily causal. It could be that both changes are independent responses to the environmental stressor of low oxygen levels and should be further investigated. When examining response time (Figure 2) and first choice (Figure 3) in relation to the mating behaviour of males from the F0 generation, it was found that the groups with lower oxygen levels (D and WD) tended to have fewer positive responses. In the subsequent generation, the association between these two behavioral features and the two oxygen-saturated groups (C and W) did not change. It is worth recalling, however, that the differences mentioned above in response time and first choice were not statistically significant. This suggests that activity rate could be a more sensitive indicator of the effects of reduced oxygen availability on chemosensory-dependent mating mechanisms of *G. locusta* than the other variables. The activity rate is directly related to the availability of oxygen in the environment. As such, it is probably more easily influenced by changes in oxygen levels, while the other variables are more complex and may be more influenced by other factors.

Overall, these results suggest that environmental stress exposure of *G. locusta* from F0 generation has a recognizable negative effect only when at least two stressors are combined. This is demonstrated in the findings of Egilsdottir et al. (2009), in which exposure to a combination of ocean acidification and decreased salinity delayed embryonic development of *Echinogammarus marinus*, but acidification alone had no effect. Similarly, Costa et al. (1998) found that, while decreased salinity had a negative effect on gammarid survival, this effect was exacerbated by the addition of increased temperature. These results suggest that it is important to consider the potential impacts of multiple environmental stressors on gammarid populations, as they could entail having a more significant effect than exposure to a single stressor. It should be noted, however, that these studies did not examine the cumulative effects of stressor exposure on subsequent generations.

In the present study, the F1 generation of *G. locusta*, whose parents have been exposed to combined stressors (i.e., WD), displayed a greater ability to cope with these environmental changes than their predecessors. F1 males exposed to WD spent more time in the association area adjacent to the female cue when compared to the previously exposed parental (F0) generation. Nonetheless, this time was not greater than that of F1 males from C or W treatment with 100% air saturation. This finding is consistent with the results of Kong et al. (2019), who found that the offspring of mussels (*Mytilus edulis*) exposed to acidification and hypoxia was less negatively affected by these stressors than the offspring of the control parents. However, as observed in both the present study and the work of Kong et al. (2019), the developmental traits of the offspring from exposed parents were still negatively affected compared to those of the control animals. This suggests that while parental exposure can provide some level of protection, it is not sufficient to completely alleviate the negative effects of environmental stressors (Kong et al. 2019). In contrast, a study on the mollusc *Saccostrea glomerata* found that a third generation exposed to high levels of CO_2_, exhibited positive carry-over effects, acquiring enhanced resistance to ocean acidification compared to the offspring with no prior history of high CO_2_ exposure (Parker et al. 2015). These results highlight the complex and potentially beneficial effects of transgenerational exposure to environmental stressors.

On the contrary, research also shows negative carry-over effects of parental stress exposure on their offspring. For instance, Borges et al. (2018a) found that during transgenerational acidification exposure, *G. locusta* parents under high CO_2_ had a significantly lower survival rate compared to the control groups. Similar negative transgenerational effects of stress exposure have been observed in fish. Wang et al. (2016) found that hypoxia caused retarded gonad development, decreased sperm count and sperm motility in the exposed parental generation of medaka fish (*Oryzias melastigma*), as well as in the F1 and F2 generations that had not been directly exposed to hypoxia. Similarly, Lai et al. (2019) demonstrated that hypoxia triggered reproductive impairment in female fish of the same species, resulting in a significant decrease in hatching success in the F2 generation. Shama et al. (2016) showed that the maternal environment had a strong influence on the phenotype of marine stickleback offspring, affecting gene expression, mitochondrial respiration, and growth. The effects of the grandmaternal environment were also observed in the F2 generation, indicating the persistence of these effects (Shama et al. 2016). To draw yet another taxon for comparison, a study by Zhao et al. (2018) showed that long-term exposure of the sea urchin *Strongylocentrotus intermedius* to high temperatures (~ 3°C above ambient) had negative transgenerational effects on hatchability and most traits of larval size. These studies suggest that the parental environment can strongly impact subsequent generations and highlight the importance of considering transgenerational effects in research on environmental stressors.

Collectively, most of these studies suggest that hypoxia, being the result of deoxygenated waters, poses a significant threat to the sustainability of natural populations of a wide variety of organisms. The present study also showed transgenerational detriments in male mating behaviour of offspring whose parents were exposed to deoxygenation, but only when reduced oxygen concentration was the only stressor. Lack of oxygen can lead to decreased energy availability and reduced metabolic rates (Borges et al., 2018b), which may explain the observed effects on mating behaviour. It is worth noting that the present study only tested one parental generation (F0) with a single subsequent generation (F1), so it is not possible to draw conclusions about the long-term effects of oxygen deficiency (OD) on transgenerational effects. To gain a more comprehensive understanding of the effects of OD on different generations and under different conditions, reciprocal transplants could be employed. This would involve exposing individuals from different generations to different treatment groups and comparing their performance. Additionally, a multigenerational approach would provide a more complete picture of the long-term impacts of OD on natural populations.

Although ocean deoxygenation is recognized as a major contributor to global biodiversity loss, its effects are largely unexplored for most marine taxa, including *G. locusta* (Borges et al., 2022). The only examination on the resistance of this species to oxygen deficiency was conducted by Bulnheim (1979), who compared a total of five *Gammarus* species, of which *G. locusta* showed the lowest survival rate when exposed to oxygen-depleted brackish water, indicating its susceptibility to low DO levels.

## 5. Concluding remarks and future research

While much progress has been made in understanding the impacts of climate change on animal behaviour, the long-term effects of transgenerational acclimation are still not fully understood. The results of this study suggest that ocean warming and ocean deoxygenation, particularly as combined multi-stressors, have transgenerational impacts on the reproductive behaviour of *G. locusta*. The observed rapid adaptability of the F1 generation may be a transgenerational effect that helps to compensate for the negative effects of deoxygenation as a single stressor in subsequent generations. Further research examining these stressors over more than just two generations could provide insights into behavioral transgenerational effects at the grand-parental level and beyond. Indeed, climate-change-related stressors only hindered population growth if the second generation was considered. Reciprocal transplants could be used to expose subsequent generations to different treatment groups and compare their response, allowing for the observation of a first exposure effect in gammarids with varying previous parental conditions. It should also be noted that the mere warming of the environment of *G. locusta* did not appear to have any significant detrimental effects on the mating behaviour of the amphipod.

There is a great lack of information on the combined effects of multiple stressors, especially on a transgenerational level. Neglecting this co-occurrence can have significant implications for research outcomes, as single stressor experiments may underestimate the consequences of, for example, ocean deoxygenation, which rarely occurs alone in marine ecosystems. Future research at the physiological and genetic levels could help shed light on the divergent behaviors of *G. locusta* under different environmental conditions.

Finally, it is worth remembering that coastal and estuarine ecosystems provide a wide range of goods and services. More research on the impacts of ocean warming and deoxygenation on various coastal organisms is needed to better understand the functioning of these ecosystems and contribute to their better management and conservation.

## Acknowledgments

We acknowledge our colleagues from Laboratório Marítimo da Guia who provided additional input to make this experiment possible.

