## Supplementary material for "Transgenerational exposure to deoxygenation and warming disrupts mate-detection in *Gammarus locusta*": Table S1

**Table S1:** Seawater physicochemical parameters in all experimental treatments. Salinity, pH, temperature, and dissolved oxygen were measured daily and averaged over the whole experimental treatment. Values for represented as mean ± standard deviation.

| | **Salinity** | **pH** | **T (°C)** | **$O_2$% air sat.** | **$O_2$mg/L** |
|---|---|---|---|---|---|
| **Control (C)** | 34.93 ± 0.4 | 8.09 ± 0.04 | 18.1 ± 0.13 | 101.59 ± 0.95 | 7.78 ± 0.16 |
| **Warming (W)** | 34.98 ± 0.58 | 8.07 ± 0.04 | 20.95 ± 0.22 | 101.48 ± 0.96 | 7.37 ± 0.08 |
| **Deoxygenation (D)** | 35.17 ± 0.42 | 8.11 ± 0.04 | 17.96 ± 0.19 | 90.67 ± 2.5 | 7.05 ± 0.47 |
| **Warming + Deoxygenation (WD)** | 35.08 ± 0.42 | 8.08 ± 0.03 | 21.05 ± 0.41 | 90.6 ± 2.62 | 6.58 ± 0.13 |
