## Supplementary material for "Transgenerational exposure to deoxygenation and warming disrupts mate-detection in *Gammarus locusta*": Figure S1

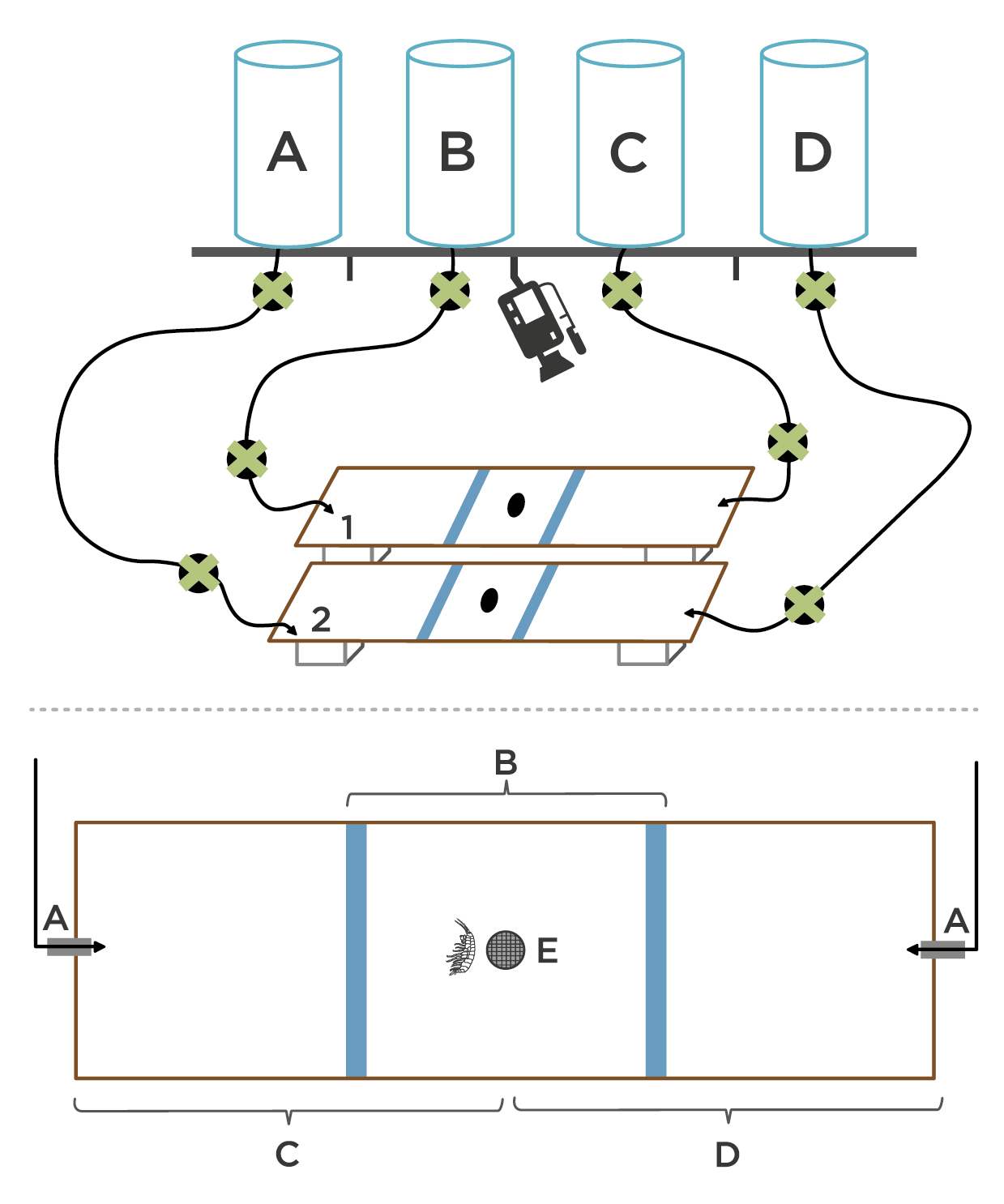

b)

a)

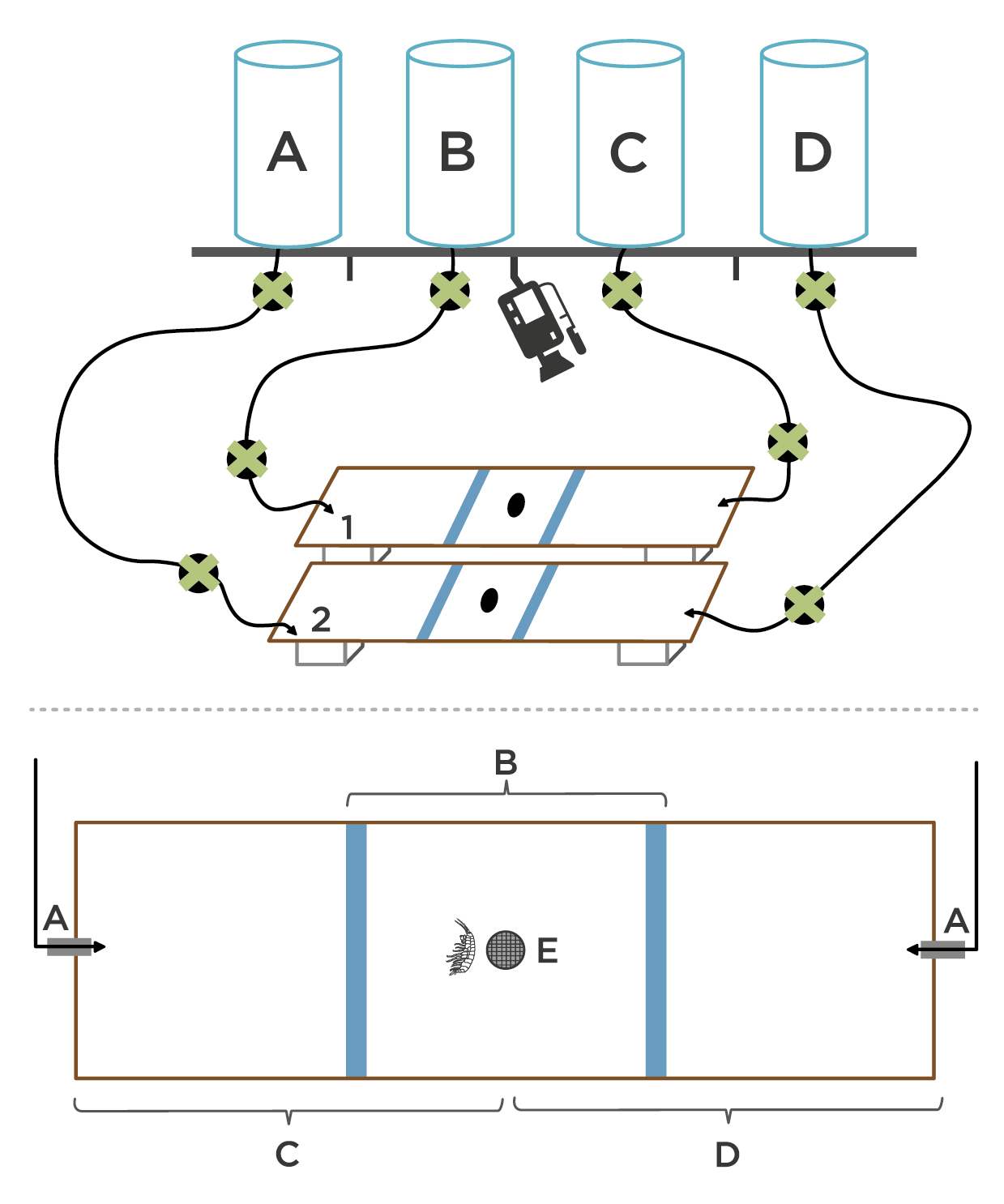

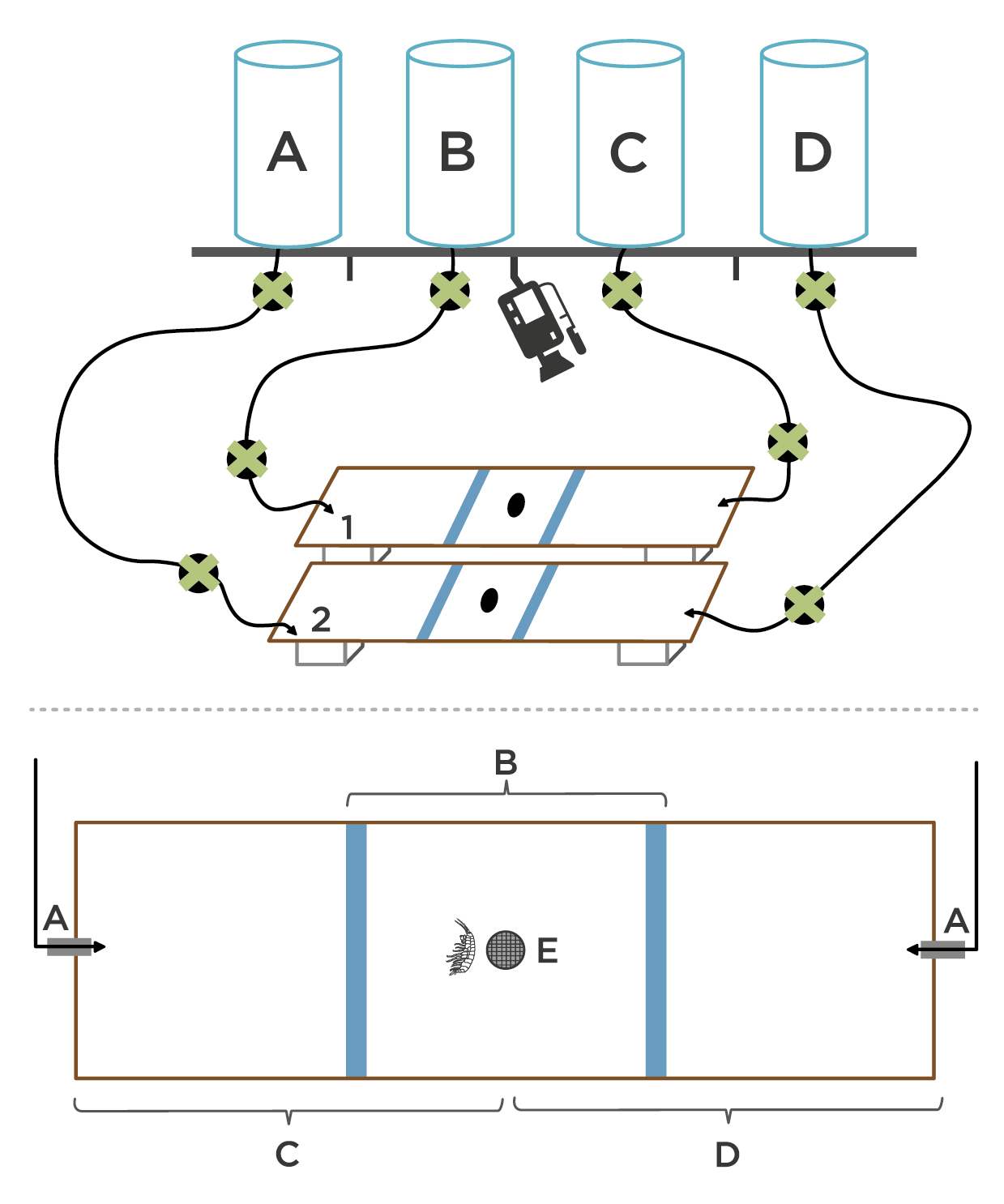

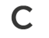

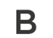

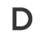

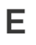

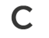

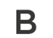

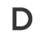

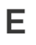

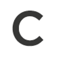

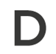

**Figure S1:** Schematic illustrations of the binary-choice set-up. a) Full experimental setup with two rectangular arenas (1&2) placed on top of two identical boxes on a leveled surface. Each arena was connected to two of four 7-L header tanks (A-D) on both sides with two distinct olfactory cues (i.e., female scent and clean seawater). The connecting tubes are all equipped with two individual valves (X) for flow-rate calibration and water-inlet control. A video camera was placed above the arenas for the recording of the test runs. b) Choice-arena. A - cue input from water tanks; B - center outlet with mesh; C - starting zone; D&E - association zones.
